# Intra-clustering analysis reveals tissue-specific mutational patterns

**DOI:** 10.1101/2024.02.26.582027

**Authors:** Stamatis Choudalakis, George Kastis, Nikolaos Dikaios

## Abstract

The identification of tissue-specific mutational patterns associated with cancer development is challenging due to the low frequency of certain mutations and the high variability among tumors within the same cancer type. To address the inter-tumoral heterogeneity issue, our study aims to uncover infrequent mutational patterns by leveraging clustering analysis. To that end, a Network Graph of 8303 patients and 198 genes was constructed from single-point-mutation data that were retrieved from The Cancer Genome Atlas (TCGA). Patient-gene groups were retrieved with the parallel use of two separate methodologies based on the: (a) Barber’s modularity index, and (b) network dynamics. An intra-clustering analysis was employed to explore the patterns within smaller patient subgroups, which involved the Fisher’s exact test, multiple correspondence analysis and DISCOVER. This analysis was applied over 24 statistically meaningful groups of 2619 patients spanning 21 cancer types and it recovered 42 mutational patterns that are not reported in the TCGA consortium analysis publications. Notably, our findings: (i) suggest that AMER1 mutations are a putative separative element between colon and rectal adenocarcinomas, (ii) highlight the significant presence of RAC1 in head and neck squamous cell carcinoma (iii) suggest that EP300 mutations in head and neck squamous cell carcinoma are irrelevant of the HPV status of the patients and (iv) show that mutational-based clusters can contain patients with contrasting genetic alterations.

**Significance:**

- The submitted paper proposes a novel method to identify infrequent cancer-specific mutational patterns through network graphs and clustering analysis.
- This analysis recovered: (a) 3 statistically significant cancer-gene relations that are not reported in the corresponding TCGA consortium analyses, (b) 28 known cancer-gene relations, (c) 39 significant patterns of co-occurrence or of mutual exclusivity that are not reported in current literature and (d) 14 reported significant mutational patterns.

## 1 Introduction

Cases of Unknown Primary (CUP) refer to aggressive cancers that have metastized to organs, and the origin of the primary tumor is unknown [1]. These cases represent approximately 3 − 5% of all cancer types, and they generally have low survival rates [1]. Knowledge of the primary cancer site in CUP cases is essential for the use of site-specific treatments and the development of suitable therapeutic plans [1, 2]. To determine the primary cancer site, physical exams and a number of lab tests and scans, such as the PET-CT scan, are conducted [3]. Despite the abundance of such procedures, there are cases where the site of origin cannot be determined, even after autopsy [4].

Recent studies have focused on the use of molecular data to define the primary cancer site in cases of unknown primary [1, 2]. Systematic analysis of gene expression, DNA methylation, mRNA expression, miRNA expression, and protein expression data have revealed distinct subtypes across cancer types [5]. However, these subtypes are not cancer-specific in a sense that samples from various cancer types might coexist in a subtype. Furthermore, these molecular data can be affected by factors other than cancer [6]. On the other hand, somatic mutations are proved to be a reliable and stable source of data [6]. For that reason, tumor prediction models mostly rely on genomic abberations such as point mutations, mutation rates, and copy number alterations [7, 8]. Nonetheless, only a small fraction of mutations may be able to confer to cancer [9]. Mutations that initiate and/or promote tumorigenesis are called driver mutations, while the genes that harbor such mutations are called driver genes. The rest of the mutations are called passenger mutations [9].

Somatic mutations can be studied based on the cell-of-origin or based on the primary cancer site. Various mutations that affect the same target cell result in different tumor types, causing inter-tumoral heterogeneity between different cancer types. Mutational differences between cancers are not only limited to the disease site. Certain cancer types may be divided into subtypes, according to the genetic alterations of cells in the tissue that account as cell-of-origin [10]. Diverse subgroups are observed even within the subtype itself [11]. Intra-tumoral heterogeneity is also present, as only a fraction of the tumor’s cells will be mutated. Other reasons causing intra-tumoral heterogeneity include the contamination by normal cells [12].

Consequently, data of somatic mutations in tumor samples are very sparse [13]. This high heterogeneity makes it difficult to recognize tissue-specific mutational patterns, as it reduces the levels of the desired signal from driver mutations. The presence of both driver and passenger mutations in the genome further reduces signal-to-noise ratio [14], while it is known that most driver genes are mutated at intermediate or low frequencies [15]. In addition, the mutational profile of patients with tumors from the same cancer subtype rarely has the same alterations [16].

Studies combining somatic mutations along with other data have revealed subtypes that, in general, include samples of various cancers per cohort [5, 14]. As pointed out in [5], the cell-of-origin may not fully determine tumor classification, but even so, it influences it. Therefore, other studies have also focused on subtype identification with a priori knowledge of the primary site [6, 17].

The applications of the analysis of somatic alterations with respect to the disease site also extend to a rapid diagnosis and treatment through the identification of disease-specific mutations found in cell-free circulating tumor DNA (ctDNA) and circulating tumor cells (CTCs) [18] through blood or urine sequencing [8].

To perform genomic-based cancer diagnosis several studies have focused on the development of network-graph-based methods [16, 19]. One reason to explain this scientific trend is that biological interactions can be interpreted as a network. The structural elements required to build a network are: (a) the set of elements-of-study (nodes) and (b) the relation(s) between these elements (edges). Therefore, any pairwise interaction can be expressed as a network. Such a network can also be presented as a graph, allowing the use of graph-theoretic tools to provide insights into the network.

In this project, we seek to recover distinct mutational patterns that are present at intermediate or low frequencies and that are able to differentiate the tissue-of-origin, under a network-graph embedding. Given that the number of patients of a specific cancer type that exhibit these patterns is expected to be very small, we focus on finding groups of patients with similar genetic traits over a suitable network, regardless of the group size. Clustering provides the most appropriate workspace of this task, as it collects the most similar nodes based on their relation, where the size of the resulting group is not necessarily fixed.

To assess the potential of clustering in the identification of infrequent mutational patterns, we constructed a Network Graph using data of high-impact somatic-point mutations over a set of 198 known driver genes and 8303 patients. Subsequently, in order to enhance accuracy, we employed two cutting-edge clustering analysis pipelines on this network, each varying in nature, and conducted a rigorous statistical analysis over the final partitions. Our analysis employed intra-clustering analysis using a combination of the Fisher’s exact test, multiple correspondence analysis and DISCOVER [20], to assess specific cancer-gene intra-cluster relations, as well as patterns of mutual exclusivity, which are known to be implicated in carcinogenesis. The clustering analysis pipelines and the statistical analysis are carefully designed in order to avoid known methodological mistakes in clustering biological data [21].

## 2 Materials and methods

### 2.1 Data Acquisition

In order to construct a gene-patient network, data from the TCGA-PanCancer Atlas project were downloaded from cBioPortal [22, 23], encompassing the thirty cancer types detailed in the results. Two of these cancer types include colon adenocarcinoma and rectal adenocarcinoma. Patients of both cancer types were included in a single dataset (colorectal adenocarcinoma), along with mucinous adenocarcinoma of the colon and rectum patients. Patients of this mixed cancer type were excluded from the network.

### 2.2 Data Preprocessing

To minimize noise, hypermutated samples (those with over 3000 mutations across all genes) were excluded from the dataset. After this filtering, mutations in 198 known cancer driver genes (whole exome sequencing), as derived from [24], were included in the curated dataset. Only high-impact mutations, including missense, nonsense, nonstop, frameshifts, in-frame deletions and insertions, and mutations occurring in splice sites [19], were considered. Metastatic samples were also removed to enhance data homogeneity.

### 2.3 Network Creation

The curated dataset was utilized to construct a network representing 30 cancer types, based solely on somatic point mutations. Each edge in the network represents a mutation in the patient on one endpoint, harbored in the gene of the other endpoint. In that context, the network is bipartite as existence of patient-to-patient or gene-to-gene edges in this network will have no physical meaning. The bipartite setting of the network allows for the utilization of tools specifically designed for this topology [25, 26].

Various tools and metrics are available for the characterization of the networks. One such tool, applicable to bipartite networks, is the biadjacency matrix. This matrix is of size *m* × *n*, where *m* and *n* represent the number of nodes in the two sets that constitute the network. In this context, *m* denotes the number of genes, while *n* corresponds to the number of patients. Another metric employed is the network density. To calculate the density of a bipartite network, the number of the edges *E* in the network is compared to the number of edges in a fully connected bipartite network with an equivalent number of nodes, distributed across the two sets that constitute the network. The maximum number of edges is given by the combinations between the two disjoint sets, i.e., *m* × *n*. The density in bipartite networks can be defined mathematically as: *D* = *E*/(*m* × *n*).

In this context, the biadjacency matrix {*a*_*ij*_} of the network is binary, with “1” indicating the presence of at least one of the aforementioned mutations in gene *i* of patient *j*, while “0” denotes the absence of such mutations. The size of the biadjacency matrix of the network was 198 × 8303 (198 genes and 8303 patients) containing 34994 edges. This yields a density of approximately 0.0213, leading to the deduction that the network is highly sparse.

### 2.4 Methods Overview

The methodology employed to unveil the underlying structure of the network involved four algorithms: Simulated Annealing (SA), Unweighted Pair Group Method with Arithmetic Mean (UPGMA), Infomap, and Order Statistics Local Optimization Method 2 (OSLOM2). The methodology employed to recover statistically significant patterns involved two algorithms: Multiple Correspondence Analysis (MCA) and Discrete Independence Statistic Controlling for Observations with Varying Event Rates (DISCOVER).

SA was implemented through the MODULAR software [25], which provides algorithms that aim to maximize the modularity index. In bipartite networks, the Barber’s modularity index [26] is commonly utilized, serving as the reference index of this study. The index can be mathematically calculated by Eq (1):

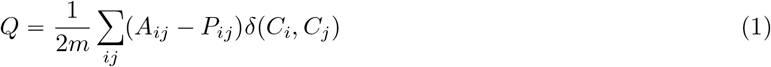

where m is the number of edges of the network, *A*_*ij*_ is the adjacency matrix of the network, *P*_*ij*_ is the adjacency matrix of the null model, and *δ* is the Kronecker delta for *C*_*i*_, *C*_*j*_ clusters. Eq (1) reveals that algorithms based on modularity maximization, rely on simple edges between the nodes of the network. A higher index value, approaching 1, signifies a more favorable corresponding partition. Since modularity maximization is an NP-hard problem, SA approximates the global maximum. The resulting partition corresponds to the local maximum of the modularity index found by SA.

UPGMA, an agglomerative hierarchical clustering method, based on the average dissimilarity index [27]:

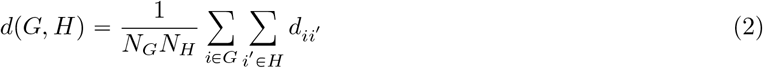

between the two clusters *G, H*, exploiting the distance matrix *D*_*ij*_ [27]. The terms *N*_*G*_, *N*_*H*_ correspond to the number of the nodes of the clusters *G, H* accordingly. The distance matrix *D*_*ij*_ has a size of (*N*_*G*_ + *N*_*H*_) × (*N*_*G*_ + *N*_*H*_), with elements *d*_*ii*_′ which correspond to the minimum distance of node *i* to *i*′. UPGMA was implemented using the “scipy.cluster.hierarchy.linkage” function of package “scipy” in Python, with the “method” parameter set to “average” [28].

Infomap [29], relying on network dynamics and minimum description length statistics (MDL), leverages random walks to cluster the network based on walk trajectories. For a given a cluster, the algorithm calculates the probability of a random walker remaining within the cluster and the probability of the random walker exiting the cluster. This information can be encapsulated in the “Map Equation”, and the resulting partition is the one that minimizes this equation. Mathematically, the map equation is expressed as:

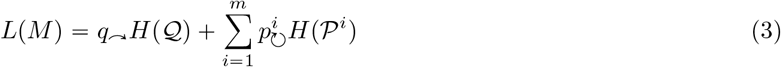

The first term of Eq (3) corresponds to the average number of bits used to describe inter-modular movements, as 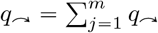 is the probability of the random walker to switch modules (per-step probability). The second term is indicative of the intra-modular movements, as 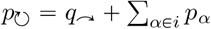., represents the number of times the codebook of module *i* was used. The 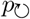 consists of the sum of the steady state distribution for all *α* nodes within the modules and the probability of exiting the module. The quantity *H*(*X*) = − ∑*p*_*i*_*log*_2_(*p*_*i*_) is the entropy of the random variable *X* with *n* states and *p*_*i*_ frequencies. A logarithm of base 2 (*log*_2_) is employed since the code lengths are measured in bits and are therefore binary.

OSLOM2 [30], a statistics-centric algorithm, aims to recover statistically meaningful modules. OSLOM2 employs a method called “Single Cluster Analysis” (SCA), which for a given *P* − *value* examines the statistical significance of each cluster locally, by iteratively adding and removing nodes. In order to recover a partition of the network, SCA is implemented multiple times. OSLOM2 also assesses if the union of two or more clusters is more statistically significant than the clusters individually, resulting in a partition with statistically significant clusters and potentially with nodes that do not belong in any cluster (homeless nodes).

Correspondence Analysis (CA) [31] is an unsupervised machine learning method of dimensionality reduction (feature extraction) of a categorical variable. MCA is an extension of Correspondence Analysis (CA) for several categorical variables, where CA is implemented on the indicator matrix (binary) of the initial data. The algorithm is based on singular value decomposition, in order to project observations and features in the low-dimensional subspace. MCA was implemented using the ‘MCA’ class of package ‘prince’ in Python [32].

DISCOVER [20] is a statistical test specifically designed to detect patterns of co-occurrence and of mutual exclusivity. The basis of this algorithm is the Poisson-Binomial distribution which does not assume that the alterations of the genes are independent and identically distributed, making it suitable for tumor-specific alteration rates. To retrieve these rates, DISCOVER solves a constrained optimization problem.

### 2.5 Clustering Pipelines

#### 2.5.1 Infomap-OSLOM2 pipeline

Infomap was utilized to establish an initial partition. Given the stochastic nature of the algorithm, Infomap underwent implementation 200 times and the outcomes were combined using Monti’s consensus clustering algorithm (see section 2.5.3). Infomap was set to recover a “two-level” partition over an “undirected” graph using the “skip-adjust-bipartite-flow” option. The consensus matrix was weighted with elements varying between 0 and 1. Any elements with a value below 0.4 [19] were zeroed, returning an unweighted (binary) final matrix 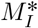. Infomap was implemented again once over the matrix 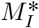, with the same parameters used in the 200 executions to get the partition *C*_*info*_.

OSLOM2 was then implemented in the partition *C*_*info*_ excluding clusters that lacked statistical significance with a P-value lower than 0.05 [19]. This led to the final partition 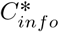. The term ‘Infomap pipeline’ will be used to denote this entire pipeline.

#### 2.5.2 MODULAR-UPGMA-OSLOM2 pipeline

The SA algorithm of MODULAR was employed to obtain an initial partition. Given the stochastic nature of the algorithm, MODULAR underwent 200 implementations and the outcomes were combined using Monti’s consensus clustering algorithm (see section 2.5.3). To address the resolution limit of modularity-based methods, a hierarchical clustering approach was utilized [33]. This limit stipulates that the resultant partition from such methods depends on the size of the network, potentially impeding the identification of small-scale modules. The application of a hierarchical algorithm to a dissimilarity matrix of the network mitigates this challenge. The consensus clustering matrix 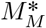 is a similarity matrix [34] of the network, and therefore, 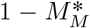 is a dissimilarity matrix (please note that 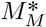 is weighted) [35]. UPGMA was then applied over 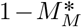, generating a dendrogram with several branches. For each branch, nodes below that branch were assigned in modules via the UPGMA, while where the rest of the nodes remained unassigned. Each unassigned node would consist of a single-node module. For every possible branch, the Barber’s modularity index (with respect to the original network) was calculated. The branch, whose corresponding partition maximized the index was considered as the optimal branch. This partition will be referred to as *C*_*mod*_.

OSLOM2 was then implemented in the partition *C*_*mod*_ partition, mirroring the Infomap pipeline, to yield the final partition 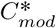. The term ‘MODULAR pipeline’ will be used to denote this entire pipeline.

#### 2.5.3 Ensemble and Consensus Clustering

Monti’s consensus clustering algorithm was employed to integrate multiple individual outcomes [35]. For each partition *h* recovered by any of the algorithms, the matrix *M*^*h*^ was established. The pair (*i, j*) in the matrix *M*^*h*^ is set to “1” if nodes *i* and *j* belonged to the same cluster in the partition *h*; otherwise the pair (*i, j*) was set to “0”. The final consensus matrix, denoted as ∑_*h*_ *M*^*h*^/*H*, was then defined, where *H* represents the total number of partitions obtained by the algorithm. A flow diagram of the data preprocessing, the network creation and the implementation of the clustering pipelines is illustrated in Fig 1.

**Figure 1.**
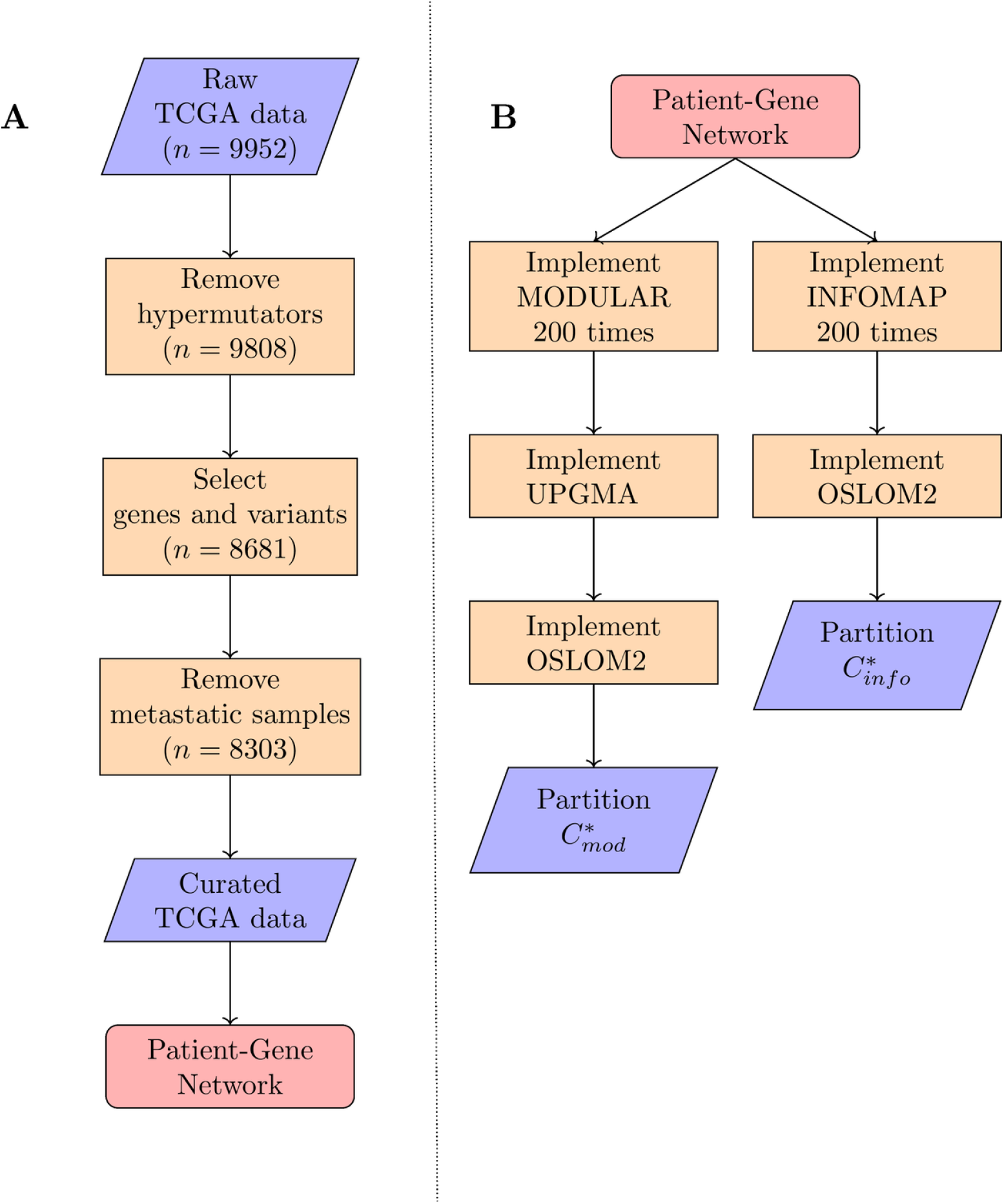
Data Preprocessing and Clustering Analysis flow diagrams. Two flow diagrams representing: (A) the data preprocessing as described in section 2.2 and (B) the implementation of the clustering analysis pipelines as described in section 2.5. In flow chart (A) the number *n* indicates the number of patients after the corresponding procedure. The flow chart (B) is separated in two columns, indicating that the pipelines where implemented independently and in parallel.

### 2.6 Threshold Filtering

The final 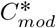 and 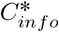 partitions contained modules with patient nodes from several cancer types. To enhance the robustness of the results, a threshold of 5% [19] per cancer type was maintained as the minimum percentage regarded significant for the outcomes of the clustering analysis pipelines. It’s worth noting that Infomap is less affected by the resolution limit compared to modularity-based methods [36].

### 2.7 Statistical Analysis

Following the threshold filtering of the results described in section 2.6, each module appearing in the final results underwent further analysis. More specifically, two separate statistical pipelines were executed for each module recovered by any of the pipelines: (i) to evaluate the significance of an intra-cluster cancer-gene relation and (ii) to assess the significance of a gene-to-gene pattern for any of the cancer types within the module.

For case (i), right-tailed Fisher’s exact tests were employed to compare the significance of the mutational load of a gene among patients of a specific cancer type with respect to the rest of the patients from other cancer types within the module. For case (ii), we implemented a combination of MCA and DISCOVER. More specifically:

- case (i): let *C* be a module with patients of the cancer types *c*_*i*_, *i* ∈ 1, 2, … *m* and genes *g*_*j*_, *j* ∈ 1, 2, … *n*. Then, *m* × *n* p-values were obtained for every cluster, using the Fisher’s exact test. To minimize the number of false positives (i.e. the number of statistically significant cancer-gene relations that are not truly significant), we implemented the false discovery rate method of Benjamini and Hochberg [37] to adjust the p-values. We set the false discovery rate threshold to be 0.05.
- case (ii): this case is based on the hypothesis that patients within the same module, have similar genetic profiles with respect to the genes of the cohort, and that patients of the same cancer type should be even more similar. Therefore, DISCOVER was run in order to obtain the initial Poisson-Binomial model of the mutational background of the patients of the cluster, regardless of the cancer type. Then, for every cancer type, MCA was performed to further reduce the number of genes. That is, for every cancer type within the module, the number of dimensions of MCA was set to be the first to exceed 99% of total explained inertia (variance). Then, following related work on MCA (package “FactoInvestigate” in R), we model the contribution of every dimension as a Gaussian, and remove patients with absolute z value of more than 1.96 (critical value of a 5% two-tailed z-test) in at least one dimension. After removing those patients, the genes with no mutation over the patients of the cluster were removed. The implementation of MCA was crucial, because it is an unbiased method to reduce false positive relations of mutual exclusivity that would occur due to the presence of genes that were mutated over a very small amount of patients of the same cancer type (e.g. one or two patients). Finally, a co-occurrence and a mutual exclusivity test were run in this curated dataset with DISCOVER, using the probabilities that were calculated across the patients of the module, regardless of the cancer type. We note that the co-occurrence tests were run only for the sake of completeness, because it is expected that patients of the same cluster, will most likely carry the same alterations over a specified set of genes, whereas the mutual exclusivity patterns are not expected through clustering analysis.

A schematic overview of our method is illustrated in Fig 2.

**Figure 2.**
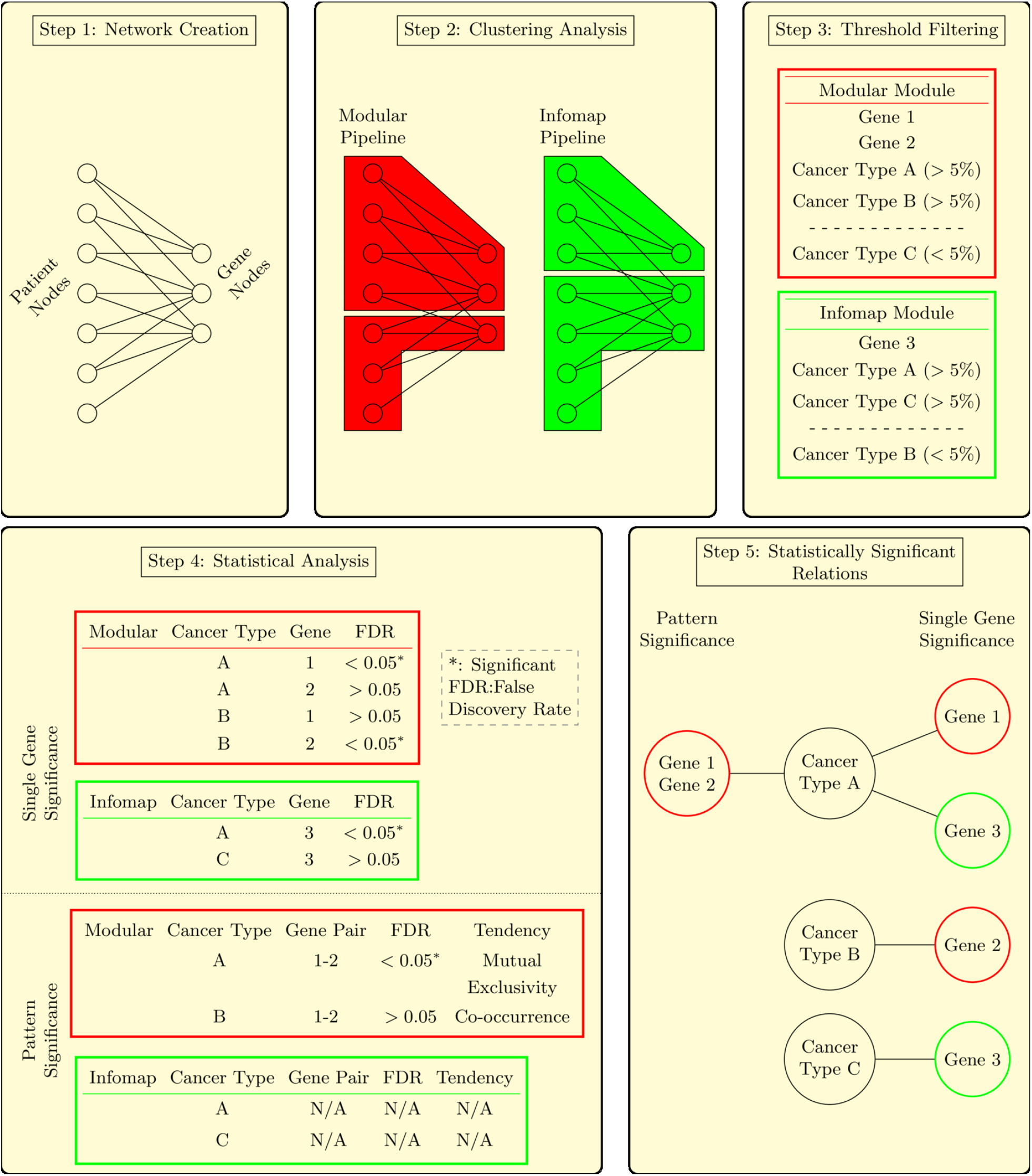
A schematic overview of the proposed method for a network with patients from cancer types A, B and C, and genes 1, 2 and 3. Step 1: The network was created from somatic-point-mutation data and is, by default, bipartite. Step 2: We use the two clustering analysis pipelines to obtain two different partitions of the network in clusters. Step 3: We filtered each cluster with respect to the amount of patients per cancer type, in order to provide robust results per cancer type. Step 4: We performed Fisher’s exact test on each cluster for every cancer-gene combination in the cluster and a combination of multiple correspondence analysis and DISCOVER for every gene-to-gene pattern for every cancer type within the cluster to obtain the p-values. The Benjamini-Hochberg false discovery rate method was employed on each cluster to obtain the adjusted p-values. A relation was considered significant if the adjusted p-value was lower than 0.05. The cases of the single gene significance and pattern significance were treated separately. Step 5: Visualization of the significant relations per cancer type.

## 3 Results

### 3.1 Implementation of the clustering pipelines

An initial estimation of the modular structure of the network was assessed with respect to the two null models provided by MODULAR. The dissociation between the modularities of the null models to the network was apparent (Fig 3C) and the p-value using Welch’s t-test was close to zero in each case. Additionally, we observed that 75% of the patient nodes in the network had a degree of less or equal to 5. This indicates that, for most patient nodes, only a small amount of information contributes to the overall network magnitude.

**Figure 3.**
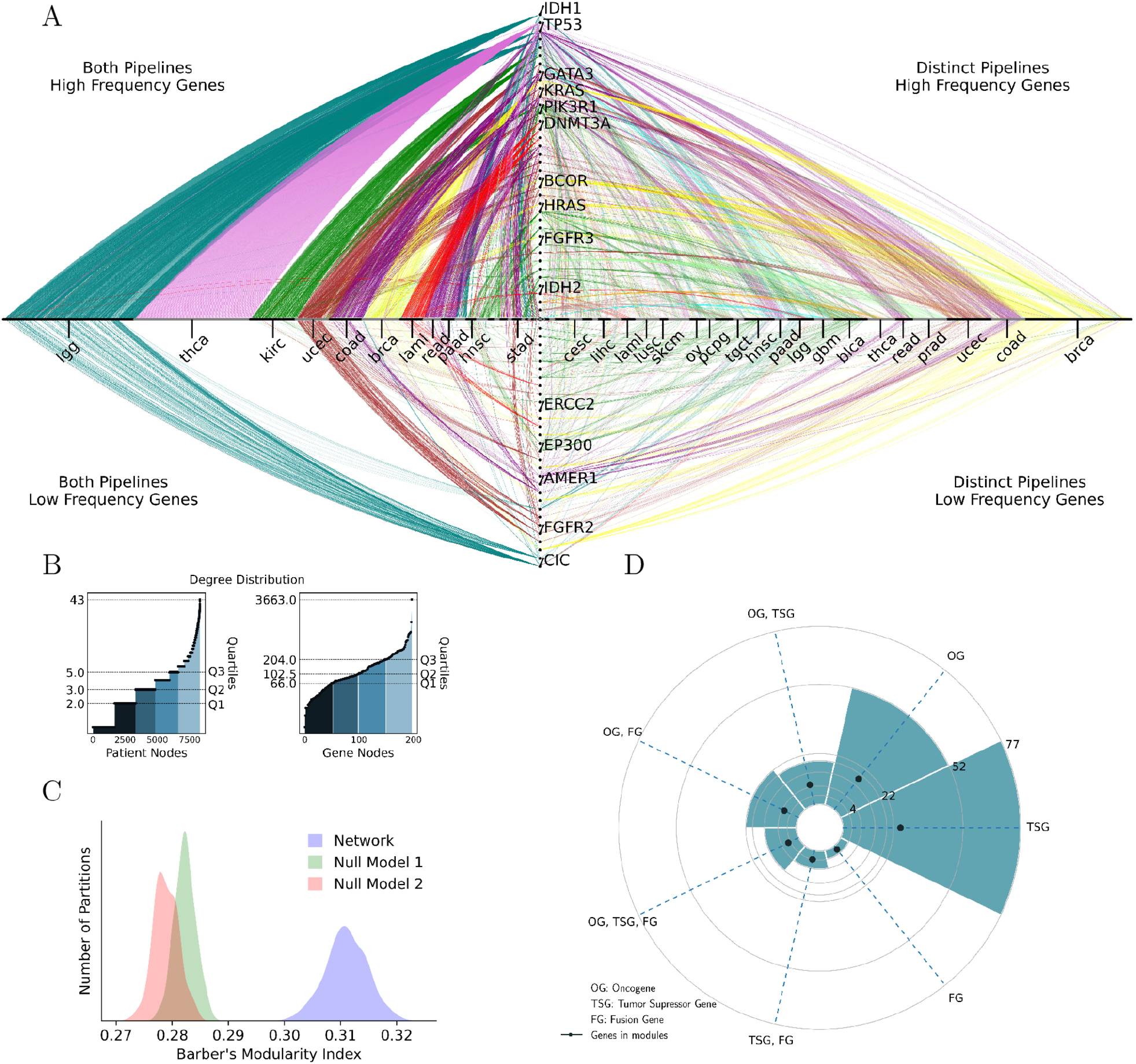
Network and node analysis. (A) A view of the network in unified clusters (See Supplementary Information for details on the unified clusters). The x-axis contains only patients that are clustered from any pipeline. If a patient is clustered from both pipelines, the corresponding node is presented at the left side of the axis. On the other hand if a patient is clustered from a single pipeline, the node is presented at the right side of the axis. The y-axis contains the genes that are assigned in modules. If a gene is mutated in a percentage higher than 30%, in any of the obtained modules, then the corresponding gene node is presented at the upper side of the axis. On the other hand, if the gene is mutated in a percentage lower than 30%, the gene node is presented in the lower side of the axis. (B) The degree distribution of the patient nodes (left) and of the gene nodes (right). The different colors of the shaded area are defined by the known quartiles. Nodes of the same shade have a degree lower than the degree that corresponds to the 25%, 50%, 75% and 100% of the degrees. For example, the darkest blue area corresponds to the patients with a degree of less or equal than 2 (=*Q*_1_).(C) The modular structure of the network is compared to the two null models provided by the MODULAR software. The dissociation of the modularities of the null models and the network is apparent. (D) The cancer-gene classification of the 198 genes of the dataset presented as a polar barplot. The black dot corresponds to the number of genes of the corresponding classification assigned in clusters. The gene classification is described at the legend below the plot.

Despite the network’s sparsity, the clustering pipelines were implemented as described in section 2.5, and approximately a third of the nodes were assigned to meaningful clusters. Specifically, the MODULAR pipeline recovered a total of 1940 (23%) patients and 66 (33%) genes in statistically significant modules, while the Infomap pipeline recovered a total of 1925 (23%) patients and 46 (23%) genes in statistically significant modules. In summary, 2619 (32%) patients and 69 (35%) genes were assigned to statistically significant modules through the union of the two final partitions. This was the only part of the project where patients from every cancer type are considered in order to establish the modular structure of the network.

A threshold filtering for each pipeline, as described in section 2.6, was applied due to the presence of patients spanning multiple cancer types in the modules. This procedure filtered out the most prevalent cancer types in each module. Tables 1 and 2 summarize the results of the MODULAR and the Infomap pipelines, referring to 17 and 19 cancer types, respectively. The total number of distinct cancer types presented in the final partitions is 21. Detailed tables of the node assignments in clusters, containing every cancer type, are presented in the Supplementary Information.

**Table 1.**
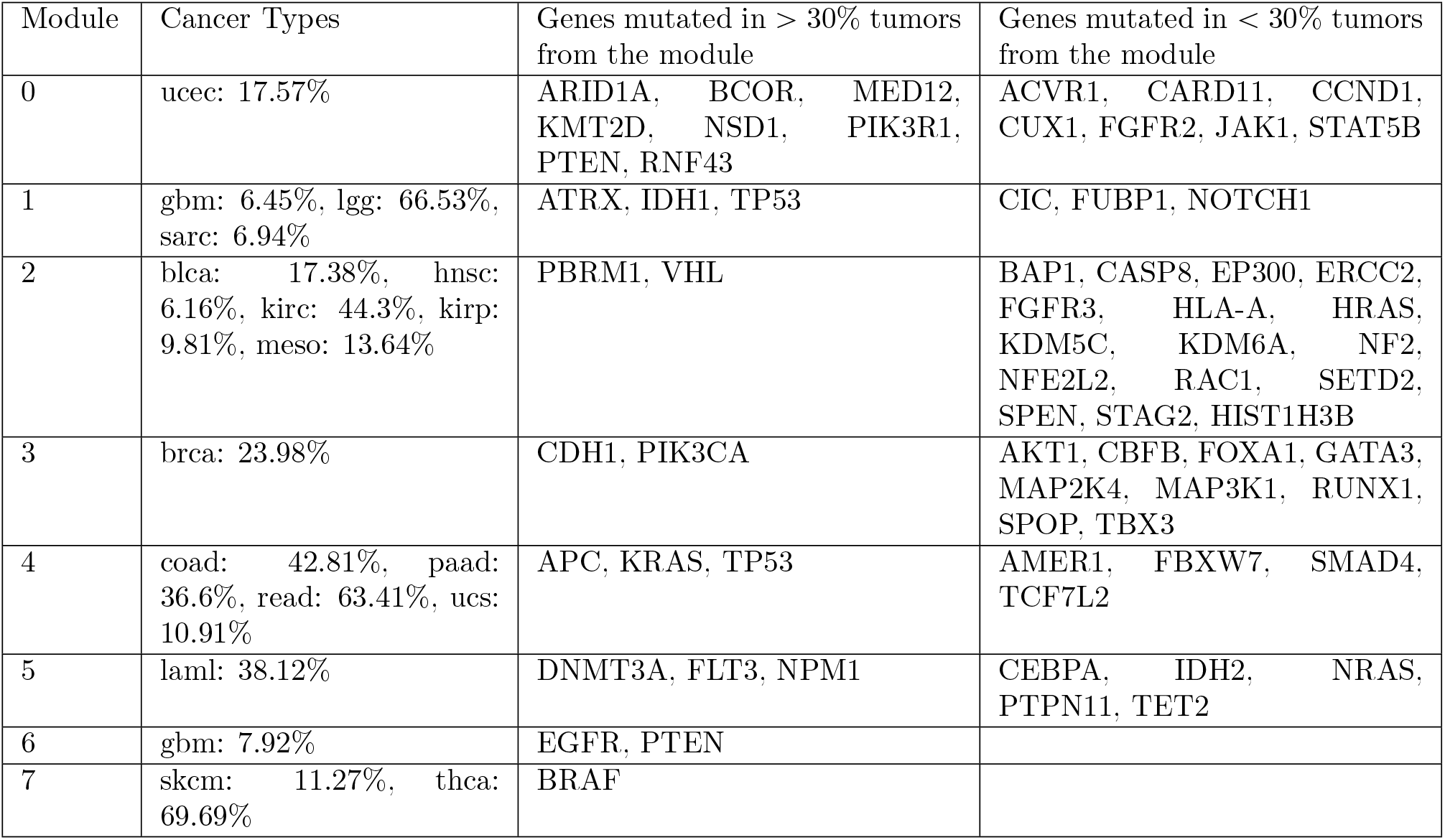
Modules recovered after ensemble clustering of 200 MODULAR partitions (consensus clustering), UPGMA and OSLOM2 implementation.

**Table 2.**
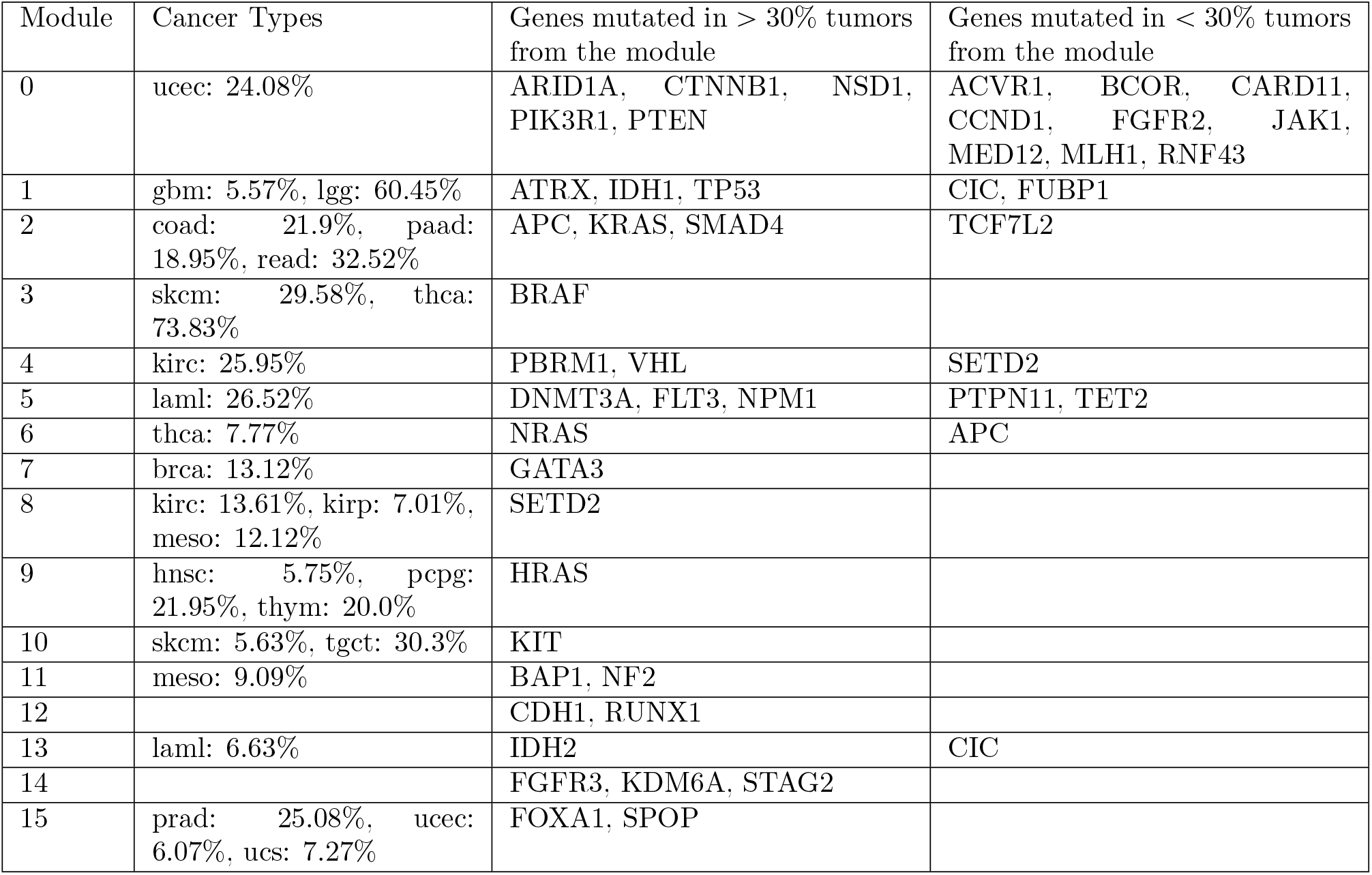
Modules recovered after ensemble clustering of 200 Infomap partitions (consensus clustering) and OSLOM2 implementation.

It should be noted that Modules “12” and “14” of the Infomap pipeline did not contain a significant number of patients in any cancer type and was therefore excluded from further investigation.

The full names of the 21 cancer types are presented in the Results section along with their TCGA abbreviations in the parentheses. The 9 cancer types not present in the final results are: Adrenocortical carcinoma (acc), Cervical squamous cell carcinoma and endocervical adenocarcinoma (cesc), Esophageal carcinoma (esca), Kidney Chromophobe (kich), Liver hepatocellular carcinoma (lihc), Lung Adenocarcinoma (luad), Lung squamous cell carcinoma (lusc), Ovarian serous cystadenocarcinoma (ov) and Stomach adenocarcinoma (stad).

An overview of the network analysis is illustrated in Fig 3.

### 3.2 Module analysis

Out of a total of 30 cancer types included in the network, 21 cancer types were significantly represented (as described in section 2.6) in modules recovered by either of the two clustering analysis pipelines. These cancer types, along with the genes that co-exist in the same module, were extensively analyzed to evaluate the performance of the pipelines in recovering statistically significant mutational patterns (see Fig 4). The results are presented below:

**Figure 4.**
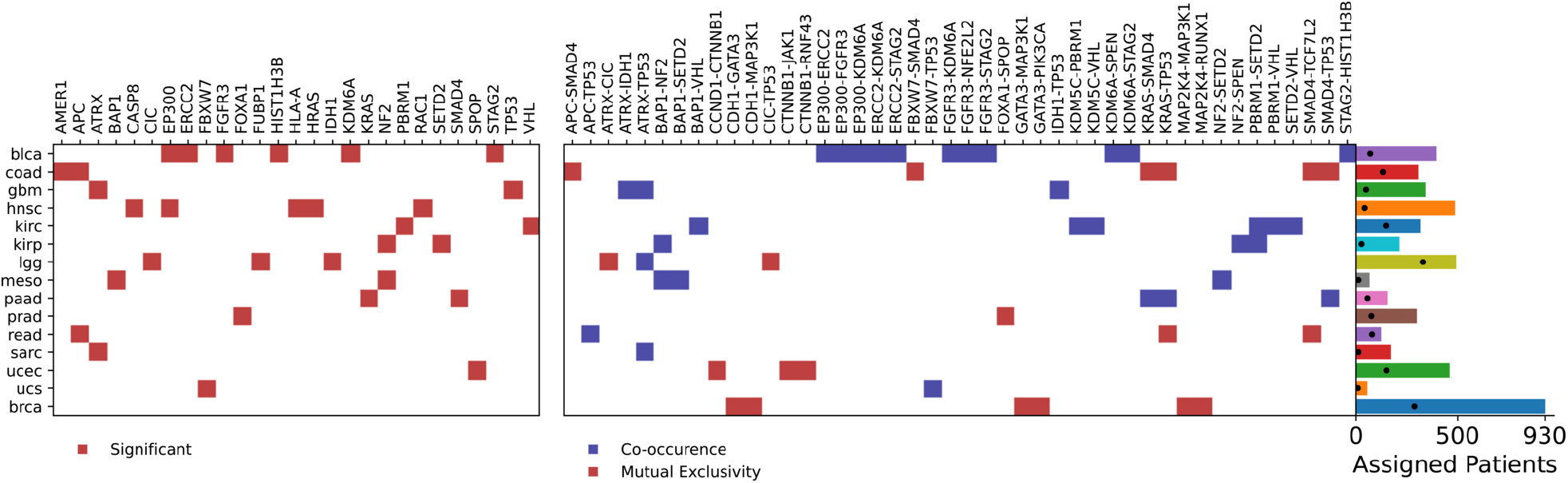
Significant relations per cancer type. This figure summarizes the significant relations and the number of clustered patients per cancer type. A significant relation is referred to as significant if the adjusted p-value from the Benjamini-Hochberg false discovery rate is lower than 0.05. At the left heatmap, the significant cancer-gene relations are marked in red. At the center, the significant gene-to-gene relations are marked in a color. If the color is red, the significant pattern is a pattern of mutual exclusivity, and if the color is blue, the significant pattern is a pattern of co-occurrence. At the right, the length of each bar of the bar plot is equivalent to the number of patients of the curated dataset of the cancer type corresponding to this bar. Furthermore, the black dot indicates the number of patients being clustered in the corresponding cancer type.

- **Bladder Urothelial Carcinoma (blca):**Eigtheen genes were clustered with 69 bladder urothelial carcinoma patients (17.38% of bladder urothelial carcinoma patients) in one module. STAG2, KDM6A, FGFR3, ERCC2, EP300 and HIST1H3B were recovered as statistically significant in the module. All genes were also reported as statistically significant in the corresponding TCGA publication, along with HRAS, CASP8 and NFE2L2, which were also clustered with bladder urothelial patients [38, 39]. Eleven statistically significant patterns of co-occurrence were also observed within one module: FGFR3-KDM6A, KDM6A-STAG2, FGFR3-STAG2, ERCC2-KDM6A, EP300-KDM6A, STAG2-HIST1H3B, KDM6A-SPEN, EP300-ERCC2, ERCC2-STAG2, FGFR3-NFE2L2, and EP300-FGFR3.
- **Breast invasive carcinoma (brca):** Eleven genes were clustered with 287 breast invasive carcinoma patients (30.86% of breast invasive carcinoma patients) in two modules. None of these genes was recovered as statistically significant in any of the modules. However, AKT1, CBFB, CDH1, FOXA1, GATA3, MAP2K4, MAP3K1, PIK3CA, RUNX1 and TBX3 were reported as driver genes in the corresponding TCGA publication [40, 41]. Six statistically significant patterns of mutual exclusivity were also observed within one module: GATA3-PIK3CA [42], CDH1-MAP3K1 [43], CDH1-GATA3 [44], MAP2K4-MAP3K1 [40], GATA3-MAP3K1 and MAP2K4-RUNX1.
- **Colon Adenocarcinoma (coad):** Seven genes were clustered with 131 colon adenocarcinoma patients (42.81% of colon adenocarcinoma patients) in two modules. APC and AMER1 were recovered as statistically significant in at least one module. APC was also reported as statistically significant in the corresponding TCGA publication, along with KRAS, TP53, SMAD4, FBXW7 and TCF7L2, which were also clustered with colon adenocarcinoma patients [45]. However, there is evidence in the literature to support the role of AMER1 as a driver gene in colorectal adenocarcinoma [46]. It must be noted that here, AMER1 is not detected as significantly mutated for rectal adenocarcinoma, but only for colon adenocarcinoma. Six statistically significant patterns of mutual exclusivity were also observed within at least one module: SMAD4-TP53, KRAS-SMAD4, KRAS-TP53, APC-SMAD4, FBXW7-SMAD4, and SMAD4-TCF7L2.
- **Glioblastoma (gbm):** Eight genes were clustered with 49 glioblastoma patients (14.37% of glioblastoma patients) in three modules. TP53 and ATRX were recovered as statistically significant in at least one module. Both genes were also reported as statistically significant in the corresponding TCGA publications, along with EGFR, PTEN and IDH1, which were also clustered with glioblastoma patients [47, 48]. Three statistically significant patterns of co-occurrence were also observed within at least one module: ATRX-TP53 [48], ATRX-IDH1 [48], and IDH1-TP53 [48].
- **Head and Neck squamous cell carcinoma (hnsc):** Eighteen genes were clustered with 42 head and neck squamous cell carcinoma patients (8.62% of head and neck squamous cell carcinoma patients) in two modules. HRAS, HLA-A, CASP8, EP300 and RAC1 were recovered as statistically significant in one module. HRAS, HLA-A and CASP8 were also reported as statistically significant in the corresponding TCGA publication [49]. Driver RAC1 mutations have been recovered as statistically significant [50]. High EP300 mutation rates were reported in the HPV-positive subsets of the TCGA dataset and the John Hopkins University, separately [51].
- **Kidney renal clear cell carcinoma (kirc):** Eighteen genes were clustered with 146 kidney renal clear cell carcinoma patients (46.2% of kidney renal clear cell carcinoma patients) in three modules. VHL and PBRM1 were recovered as statistically significant in one module. Both of them are also reported as statistically significant in the corresponding TCGA publication, along with SETD2, BAP1, KDM5C and NFE2L2, which were also clustered with kidney renal clear cell carcinoma patients [52]. Six statistically significant patterns of co-occurrence were also observed within one module: PBRM1-VHL, PBRM1-SETD2, BAP1-VHL, SETD2-VHL, KDM5C-PBRM1, KDM5C-VHL.
- **Kidney renal papillary cell carcinoma (kirp):** Eighteen genes were clustered with 27 kidney renal papillary cell carcinoma patients (12.62% of kidney renal papillary cell carcinoma patients) in two modules. SETD2 and NF2 were recovered as statistically significant in one module. Both genes were also reported as statistically significant in the corresponding TCGA publication, along with PBRM1, BAP1, KDM6A, NFE2L2 and STAG2, which were also clustered with kidney renal papillary cell carcinoma patients [53]. Three statistically significant patterns of co-occurrence were also observed within one module: PBRM1-SETD2 [53], BAP1-NF2, and NF2-SPEN.
- **Acute Myeloid Leukemia (laml):** Nine genes were clustered with 76 acute myeloid leukemia patients (41.99% of acute myeloid leukemia patients) in three modules. None of these genes was recovered as statistically significant in any of the modules. However, DNMT3A, FLT3, NPM1, CEBPA, IDH2, NRAS, PTPN11 and TET2 were reported as statistically significant in the corresponding TCGA publication [54].
- **Lower Grade Glioma (lgg):** Six genes were clustered with 328 lower grade glioma patients (66.53% of lower grade glioma patients) in two modules. FUBP1, CIC and IDH1 were recovered as statistically significant in one module. All three genes were also reported as statistically significant in the corresponding TCGA publication, along with TP53, ATRX and NOTCH1, which were also clustered with lower grade glioma patients [55]. Two statistically significant patterns of mutual exclusivity were also observed within at least one module: ATRX-CIC [56], CIC-TP53 [56]. One statistically significant pattern of co-occurrence was also observed within at least one module: ATRX-TP53 [56]
- **Mesothelioma (meso):** Eighteen genes were clustered with 12 mesothelioma patients (18.18% of mesothelioma patients) in three modules. NF2 and BAP1 were recovered as statistically significant in one module. Both genes were also reported as statistically significant in the corresponding TCGA publication, along with SETD2 that was also clustered with mesothelioma patients [57]. Three statistically significant patterns of co-occurrence were also observed within at least one module: BAP1-NF2 [57], BAP1-SETD2 [57], and NF2-SETD2.
- **Pancreatic Adenocarcinoma (paad):** Seven genes were clustered with 56 pancreatic adenocarcinoma patients (36.6% of pancreatic adenocarcinoma patients) in two modules. KRAS and SMAD4 were recovered as statistically significant in at least one module. Both genes were also reported as statistically significant in the corresponding TCGA publication, along with TP53 that was also clustered with pancreatic adenocarcinoma patients [58]. Three statistically significant patterns of co-occurrence were also observed within at least one module: KRAS-SMAD4, KRAS-TP53, SMAD4-TP53.
- **Pheochromocytoma and Paraganglioma (pcpg):** HRAS was clustered with 18 pheochromocytoma and paraganglioma patients (21.95% of pheochromocytoma and paraganglioma patients) in one module. HRAS was not recovered as statistically significant in the module. However, it was reported as statistically significant in the corresponding TCGA publication [59].
- **Prostate Adenocarcinoma (prad):** FOXA1 and SPOP were clustered with 75 prostate adenocarcinoma patients (25.08% of prostate adenocarcinoma patients) in one module. FOXA1 was recovered as statistically significant in one module. Both genes were reported as statistically significant in the corresponding TCGA publication [60]. One statistically significant pattern of mutual exclusivity was also observed within one module: FOXA1-SPOP [60].
- **Rectal Adenocarcinoma (read):** Seven genes were clustered with 78 rectal adenocarcinoma patients (63.41% of rectal adenocarcinoma patients) in two modules. APC was recovered as statistically significant in both modules. APC was also reported as statistically significant in the corresponding TCGA publication, along with KRAS, TP53, SMAD4, FBXW7 and TCF7L2, which were also clustered with rectal adenocarcinoma patients [45]. Two statistically significant patterns of mutual exclusivity were also observed within one module: KRAS-TP53, SMAD4-TCF7L2. One statistically significant pattern of co-occurrence was also observed within one module: APC-TP53.
- **Sarcoma (sarc):** Six genes were clustered with 12 sarcoma patients (6.94% of sarcoma patients) in one module. ATRX was recovered as statistically significant in the module. ATRX was also reported as statistically significant in the corresponding TCGA publication along with TP53 that was also clustered with sarcoma patients [61]. One statistically significant pattern of co-occurrence was also observed within one module: ATRX-TP53.
- **Skin Cutaneous Melanoma (skcm):** BRAF and KIT were clustered with 25 skin cutaneous melanoma patients (35.21% of skin cutaneous melanoma patients) in three modules. None of these genes was recovered as statistically significant in any of the modules. However, BRAF was reported as statistically significant in the corresponding TCGA publication [62].
- **Testicular Germ Cell Tumors (tgct):** KIT was clustered with 20 testicular germ cell tumor patients (30.3% of testicular germ cell tumor patients) in one module. KIT was not recovered as statistically significant in the module. However, it was reported as statistically significant in the corresponding TCGA publication [63].
- **Thyroid cancer (thca):** Three genes were clustered with 315 thyroid cancer patients (81.61% of thyroid cancer patients) in three modules. None of these genes was recovered as statistically significant in any of the modules. However, BRAF and NRAS, which were clustered with thyroid cancer patients, were considered as statistically significant in the corresponding TCGA publication [64].
- **Thymoma (thym):** HRAS was clustered with 10 thymoma patients (20% of thymoma patients) in one module. HRAS was not recovered as statistically significant in the module. However, it was reported as statistically significant in the corresponding TCGA publication [65].
- **Uterine Corpus Endometrial Carcinoma (ucec):** Nineteen genes were clustered with 148 uterine corpus endometrial carcinoma patients (32.1% of uterine corpus endometrial carcinoma patients) in three modules. SPOP was recovered as statistically significant in one module. SPOP was also reported as statistically significant in the corresponding TCGA publication, along with ARID1A, CTNNB1, PIK3R1, PTEN, CCND1, FGFR2, BCOR and RNF43, which were also clustered with uterine corpus endometrial carcinoma patients [66]. Three statistically significant patterns of mutual exclusivity were also observed within one module: CCND1-CTNNB1, CTNNB1-RNF43 and CTNNB1-JAK1.
- **Uterine Carcinosarcoma (ucs):** Nine genes were clustered with 9 uterine corpus endometrial carcinoma patients (16.36% of uterine carcinosarcoma patients) in two modules. FBXW7 was recovered as statistically significant in one module. FBXW7 was also reported as statistically significant in the corresponding TCGA publication along with KRAS, TP53 and SPOP, which were also clustered with uterine carcinosarcoma patients [67]. One statistically significant pattern of co-occurrence was also observed within one module: FBXW7-TP53.

## 4 Discussion

Our primary emphasis was on pinpointing tissue-specific patterns across 30 different cancer types, that occur in a relatively small amount of patients, using somatic-point-mutation data sourced from the TCGA. Our methodology involved employing intra-clustering statistical analysis, which has not been implicated in relevant works, to the best of our knowledge. We incorporated the use of multiple correspondence analysis and DISCOVER over gene-patient clusters derived from two clustering pipelines, which helped reveal intermediate- and low-frequency patterns that were hindered from the general population, as well as some known mutational patterns.

The key finding of our study is the identification of 3 mutated genes that were not recovered in the corresponding TCGA consortium analysis as statistically significant, which suggests their putative role as driver genes. Specifically, AMER1 in colon adenocarcinoma and RAC1 and EP300 in head and neck squamous cell carcinoma emerged as statistically significant. Currently, there is no evidence that AMER1 is found in colon or rectal adenocarcinoma, separately, as most genome-wide studies treat colorectal adenocarcinomas as a single and indistinguishable cancer type. Therefore, our findings suggest that AMER1 mutations may differ between colon and rectal adenocarcinomas. Chang *et al*. identified a specific driver RAC1 mutational hotspot (RAC1 A159V hotspot) for head and neck cancers, leveraging data from different sources, one of which is the TCGA dataset [50]. However, in their study, head and neck cancers consist of adenoid cystic carcinomas, head and neck squamous cell carcinomas, nasopharygeal carcinomas, and thyroid carcinomas. Here, we provide further evidence of the driver role of RAC1 specifically for head and neck squamous cell carcinomas, regardless of the position of the mutation and solely occupying TCGA data. Finally, EP300 is found to be significantly mutated in two separate HPV-positive subsets of head and neck squamous cell carcinoma patients, one of which being the TCGA dataset [51]. Here, EP300 is also recovered as statistically significant in a module, but only 1 out of the 30 patients of the module was HPV-positive, suggesting the presence of EP300 mutations in head and neck squamous cell carcinomas, regardless of the HPV status of the patients.

The second key finding was our method’s ability in detecting 7 known significant patterns of mutual exclusivity and 7 of co-occurrence, aligning with TCGA consortium analysis and other studies. Impressively, we identified 13 statistically significant patterns of mutual exclusivity and 26 of co-occurrence previously unreported in the current literature. The large number of co-occurrences is expected due to the nature of clustering analysis, as opposed to the number of mutual exclusivities. This not only suggests the ability of clustering to group patients with similar genetic alterations, but also with contrasting ones. This is also extended to discrepancies observed among patients of the same cancer type within different clusters. For example, the differentiation of EGFR-PTEN and IDH1-TP53-ATRX mutations in glioblastoma and the differentiation of BRAF mutations and NRAS mutations in thyroid cancer was depicted here and was consistent to the current literature [19, 48, 64]. We must note that the impact of these novel patterns on cancer initiation and/or progression remains unknown and falls beyond the scope of this study. We also take into consideration the work of Canisius *et al*., which states that chance explains most co-occurrence [20].

The third key finding was that the intra-clustering outlier detection implemented using MCA, was able to successfully detect the non-driver genes for every cancer type of the module, reducing the overall number of false positives. This indicates the potential of our approach to correctly identify distinctive features within a module and therefore distinctive cancer-gene relations with not necessarily only one prevalent cancer type within a module. However, this is not the case of Fisher’s exact test, where known significantly mutated genes were not always recovered, even when co-existing with patients of the corresponding cancer type. This discrepancy is observed across all instances, suggesting its limited suitability for detecting frequently mutated genes across the complete dataset.

A more technical insight arises from a comparison between our method and the work of J. Iranzo *et al*. [19], who conducted the most comparable study. The lower number of modules recovered by the MODULAR pipeline in our study, likely due to an increase in patients numbers, led us to hypothesize that Infomap could detect nested relations within the MODULAR baseline partition, due to its ability to overcome the well-known resolution limit of modularity-based algorithms. Indeed, Infomap increased the overall number of assigned patients from 1940 to 2619, providing meaningful results for pheochromocytoma and paraganglioma, prostate adenocarcinoma, testicular germ cell tumors and thymoma, which faced limitations in the MODULAR partition. Detailed analysis of discrepancies between the two pipelines is presented in the Supplementary Information.

Overall, the use of both clustering analysis pipelines has uncovered clusters containing reported and non-reported statistically significant relations. Further research on the non-reported relations is imperative to validate or invalidate their oncogenic role, due to several factors that have not been taken into consideration such as the mutation rates, the dna methylation status and the gene expression levels. In conclusion, our study advocates for the incorporation of intra-clustering statistical analysis in cancer genomics, emphasizing the potential insights gained by examining smaller patient cohorts.

### 4.1 Statistical Considerations

The statistical analysis employed in this study is partially based on the Fisher’s exact test. Seven clusters from the Infomap pipeline and four clusters from the MODULAR pipeline were found to predominantly contain a single cancer type. These clusters exhibit enrichment in cancer-specific driver relations, with only 11 out of the 45 cancer-gene combinations not reported as significant in the current literature. Even when the Fisher’s exact test failed to recover these relations by default, the pattern detection statistical analysis proved efficient, revealing significant patterns within these clusters. In all cases, at least one of the two genes implicated in the pattern is a known significantly mutated gene.

Regarding the gene-to-gene pattern detection, there were two cases where there was no need to implement MCA: (i) when only one gene was present within a module and (ii) when all the patients of a specific cancer type within a module, had exactly the same alterations over the genes of the module. In case (i), there was also no need to implement DISCOVER.

DISCOVER was employed in all cases where more than one gene were present within the module, after the implementation of MCA (if applicable) and it was able to run in all but one cases. This is the case of head and neck squamous cell carcinoma in module 2 of the Modular pipeline.

## 5 Conclusion

To conclude, we have proposed a method to gain tissue-specific insights by recovering statistically significant mutational patterns through network graphs and clustering analysis. The network was constructed through somatic-point-mutation data from TCGA and it contained 8303 patients and 198 genes. The clustering analysis pipelines used to obtain patient-gene groups were based on different algorithms. The first is based on the maximization of Barber’s modularity index, while the second is based on minimal description statistics and random walks. An ensemble clustering approach was adopted in both cases, in order to obtain more robust and stable solutions. Statistics to obtain the clusters was introduced, with the implementation of OSLOM2. The Fisher’s exact test was used within each cluster, followed by the Benjamini-Hochberg false discovery rate method to minimize the number of false positive significant relations. Furthermore, a combination of multiple correspondence analysis and DISCOVER was implemented in order to obtain patterns of co-occurrence and of mutual exclusivity. We note that to the best of our knowledge, this is the first time that this kind of intra-clustering analysis is performed. This analysis recovered: (a) 3 statistically significant cancer-gene relations that are not reported in the corresponding TCGA consortium analyses, (b) 28 known cancer-gene relations, (c) 39 significant patterns of co-occurrence or of mutual exclusivity that are not reported in current literature and (d) 14 reported significant mutational patterns.

## 6 Supplementary Information

### S1 File Supplementary Information file

This file contains a table of the TCGA abbreviations, complete tables of module assignment, a thorough comparison of the results of the clustering pipelines, and supplementary information regarding the clustering pipelines.

## 7 Acknowledgments

The authors would like to thank Dr. Jaime Iranzo for his guidance and advice on terms of network and pipeline construction. His thorough explanation on both technical and biological matters played a crucial role on the completion of this work. The results published here are in whole based upon data generated by the TCGA Research Network: https://www.cancer.gov/tcga. This work was funded by the Sectoral Development Program (OΠΣ 5223471) of the Ministry of Education, Religious Affairs and Sports, through the National Development Program (NDP) 2021-25.

